# Double dissociation of visuomotor interaction mediated by visual feedback during continuous *de novo* motor learning

**DOI:** 10.1101/2023.11.20.567820

**Authors:** Junghyun Kim, Sungbeen Park, Kwangsun Yoo, Sungshin Kim

## Abstract

While the sensorimotor cortices are central neural substrates for motor control and learning, how the interaction between their subregions with visual cortices contributes to acquiring *de novo* visuomotor skills is poorly understood. We designed a continuous visuomotor task in fMRI where participants learned an arbitrary visuomotor mapping. To dissociate motor and somatosensory cortices functions, we manipulated visual feedback of a cursor such that they learned to control using fingers under two alternating conditions: online cursor feedback is available or unavailable except when a target is reached. We found double dissociation of fMRI activity in subregions of the sensorimotor and visual cortices and their interaction, which were mediated by the visual feedback. We also found a significant reduction in functional connectivity between somatosensory cortices and early visual cortices, which was highly correlated with performance improvement. These findings support the distinct interaction between subregions of sensorimotor cortices and visual cortices while highlighting the more dominant role of somatosensory cortices over motor cortices during *de novo* motor learning.

## Introduction

Visuomotor learning involves the execution and correction of motor commands to achieve a task goal based on visual feedback, which is underpinned by adaptive interaction between sensorimotor and visual cortices^1,2^. It has been well established that visual feedback shapes visuomotor learning while modulating activities of neural substrates engaged in the visuomotor interaction^3–7^. For instance, we can manipulate the visibility of the feedback, which provides continuous movement of the end-effector or its position only at the end of movement^3,8,9^. The visibility could affect the integration of the visual feedback with other sensory feedback, such as proprioception, which is processed by the somatosensory cortices. Additionally, the visibility could distinctively modulate activity in early and late visual cortices as well as their interaction with sensorimotor cortices. Thus, manipulating visual feedback could be a promising approach to investigate distinct functional roles of the sensorimotor and visual subregions in visuomotor learning.

However, the motor and somatosensory cortices are often coactivated during motor tasks, as reported in previous fMRI studies using a similar manipulation of the visual feedback^8–11^. The coactivation would be primarily due to intricate interaction between motor and somatosensory cortices during movement^7^. The motor and somatosensory cortices are closely interconnected, playing roles in creating motor commands, anticipating sensory outcomes, and processing sensory feedback. The intertwined interaction makes it challenging to determine the extent to which motor versus somatosensory cortices contribute to motor learning^7^. Previous studies employed simple motor control tasks such as tracking a target^8^, moving a pendulum^9^, and hand-grasping task^10,11^, in which they have not investigated neural plastic changes of the sensorimotor network. Moreover, little has been explored about how the subregions of the sensorimotor network interact with those of the visual network, early and late visual cortices.

Here, we employed a delicate motor skill learning task in fMRI where participants used their fingers to control an on-screen cursor while learning an arbitrary hand-to-cursor mapping^12–14^. In contrast to typical motor adaptation and sequence learning tasks with well-defined sensory targets, our task requires exploration in a high-dimensional motor space while learning proprioceptive state, i.e., hand posture, which is mapped to a low-dimensional target space. Considering the vital role of afferent proprioception in the task, our aim was to separate its influence on motor learning from the effects of efferent motor execution. Furthermore, we sought to explore how their neural substrates, motor cortices, and somatosensory cortices differentially interact with visual cortices during motor learning. To this end, we manipulated the visibility of the visual feedback so that participants performed a motor learning task under two interleaved feedback conditions: continuous online visual feedback of the cursor movement and binary feedback only available when the cursor reached a target otherwise hidden. We first contrasted the overall fMRI activity in the motor and somatosensory cortices as well as that in the subregions of the visual cortices for the two visual feedback conditions. Then, we further explored the functional connectivity patterns between the sensorimotor and visual networks and correlated them with learning performance.

## Results

### Behavioral data analysis

Twenty-four participants completed the experiment. The main task began after the localizer session. The purpose of the localizer session was to define regions related to hand movement. In the main session, participants wore a data glove on their right hand and controlled an on-screen cursor by moving their right fingers. The cursor position was calculated from a predefined hand-to-cursor mapping, which was calibrated for individual participants (see Methods). Their goal in the experiment was to reach a target with a cursor as quickly as possible. Participants performed the task under two different conditions alternately (Fig. 1A). In the continuous visual feedback condition, continuous online visual feedback was provided as the position of an on-screen cursor. Additionally, a target appeared red when reached by a cursor. In the other binary visual feedback condition, the online cursor position was not provided, and thus, the target color, which appeared red when reached, was the only cue available on the screen. The hand-to-cursor mapping was identical in both feedback conditions.

**Figure 1.**
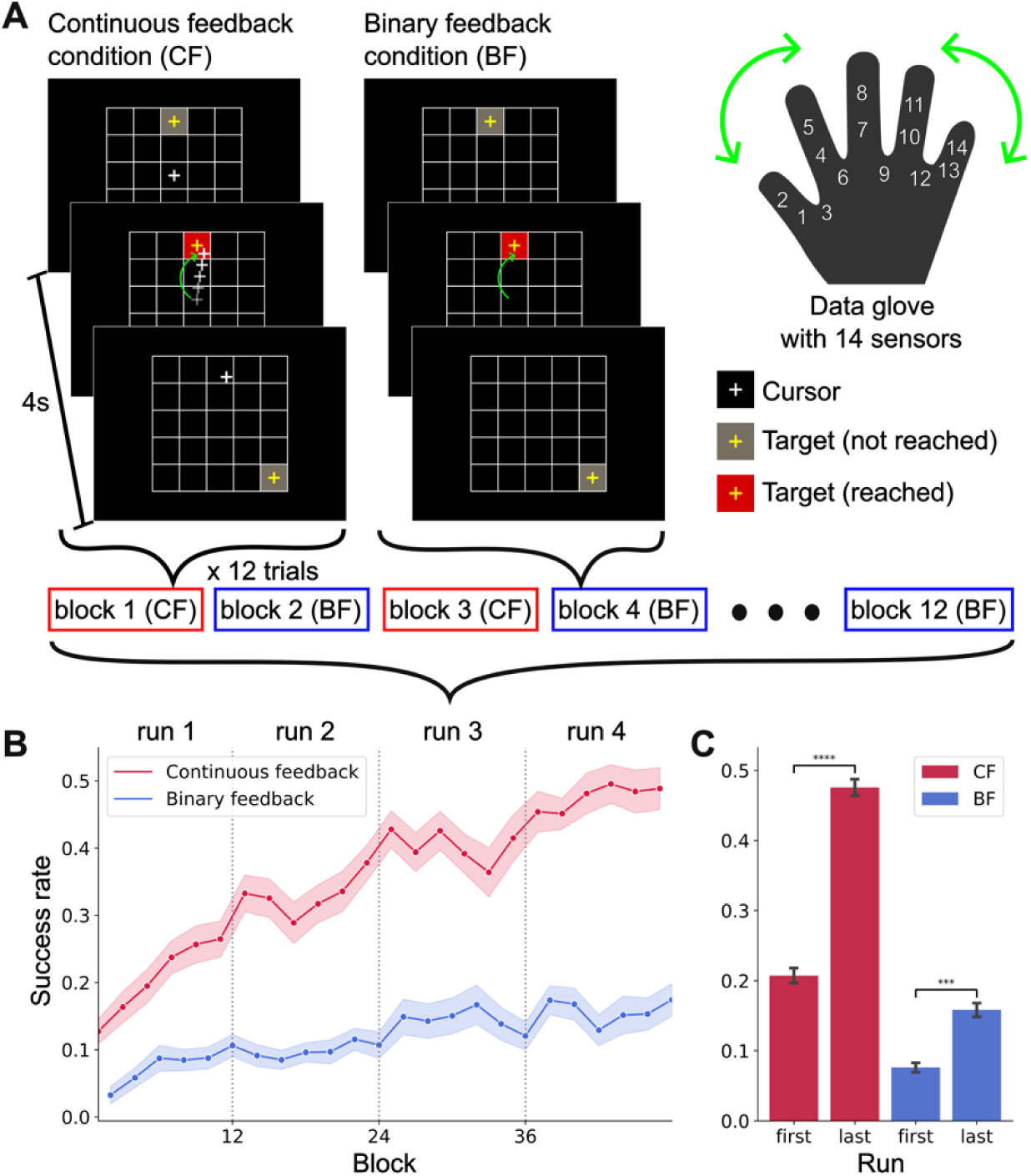
**(A)** Overview of the de novo motor learning task. Participants learned to control a cursor by moving their right fingers. Their goal was to reach a target on a 5 x 5 grid as quickly as possible and stay on the target until the next target was turned on. There were two conditions: continuous feedback (CF) and binary feedback (BF) conditions. Participants could see both cursor movement and a target in the continuous feedback condition, whereas cursor movement was hidden in the binary feedback condition. (B) The behavioral result of the task was on the continuous feedback condition and the binary feedback condition. The increase in success rate implies that participants reached the targets more quickly and stayed on them longer. Participants performed better with the continuous feedback. (C) Averaged success rates of the first and the last run from both conditions. Success rates significantly increased in both conditions, meaning they also learned the mapping in the binary feedback condition.

To quantify how much participants learned a motor skill through the experiment, we formalized success rate as a proportion of time during which a target turned red in a trial. For example, a success rate of 0.1 indicates that participants spent 3.6 seconds to reach a target and stayed in the target for 0.4 seconds. The higher the success rate indicates the faster participants reached the target, which implies participants have learned a hand-to-cursor mapping. As expected, they gradually learned the task throughout the experiment with significant improvement of the success rate from the first run to the last run both in the continuous feedback condition (*t*(23)=13.33, *p*<10^-4^) and in the binary feedback condition (*t*(23)=4.49, *p*<0.001) (Fig. 1B). The improvement of the success rate was 2 to 2.5 times higher in the continuous feedback condition than in the binary one (*t*(23)=13.97 *p*<10^-4^) (Fig. 1C), confirming that participants learned faster when online cursor feedback is provided.

### Activation in sensorimotor and visual cortices

To understand the distinct roles of sensorimotor cortices in acquiring a novel motor skill with the two visual feedback conditions, we contrasted fMRI activity between the conditions using a conventional GLM analysis. Notable, we defined regions of interest in the sensorimotor cortices on the cortical surface for all the analyses in the present study since the surface-based analysis improves the specificity of fMRI activity across folded cortical regions over the volume-based analysis^15^. In particular, the surface-based analysis is critical to dissociate fMRI activity between the primary motor (M1) and somatosensory cortices (S1), which are a few millimeters apart in 3D volume space, i.e., Euclidean space. In addition, we focused our analyses on the contralateral (left) hemisphere.

The cortical surface-based GLM analysis revealed that the dorsal premotor cortex (PMd) and higher-order visual cortices in the dorsal and the ventral visual pathways exhibited significantly higher fMRI activity in the continuous feedback condition than in the binary one. In contrast, S1 and early visual areas demonstrated significantly higher fMRI activity when the binary feedback was provided. M1 is an intermixed region where the activation levels were similar between the conditions (Fig. 2C). Notably, the activity in S1 and part of M1 is contralateral, meaning that we do not observe it on the right hemisphere. In contrast, all the other activation patterns are bilateral.

**Figure 2.**
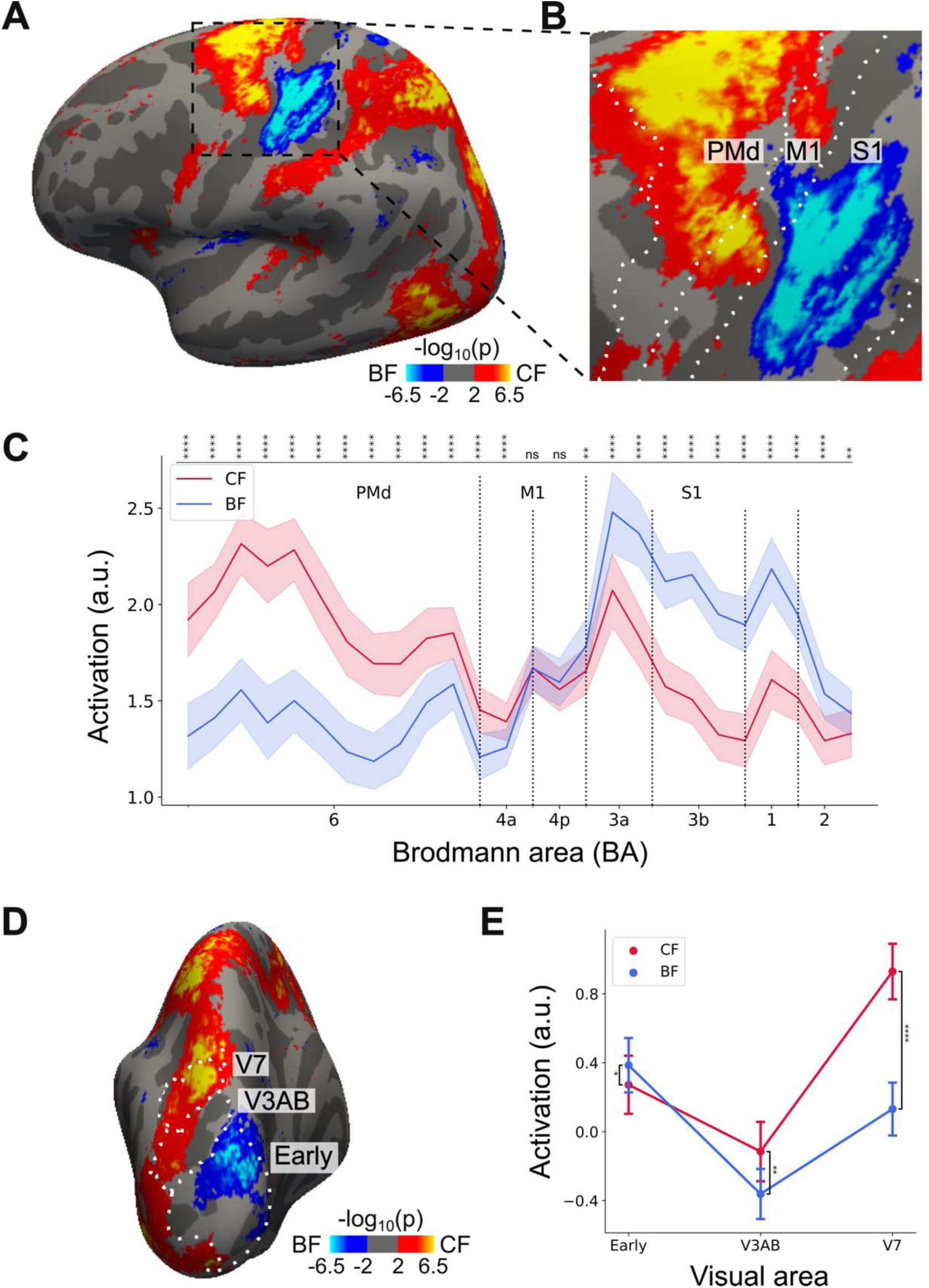
**(A)** Univariate activation map from the lateral view for the contrast continuous feedback > binary feedback. **(B)** Enlarged map for (A) highlighting sensorimotor regions. **(C)** ROI analysis (see Methods) of mean activation (± standard error of the mean). **(D)** Same as A but from the caudal view. **(E)** ROI analysis of mean activation over visual areas (± standard error of the mean). The level of significance was represented by asterisks as follows. \**p*<0.05, \*\**p*<0.01, \*\*\**p*<0.001, \*\*\*\**p*<10^-4^

To analyze the regional specificity of fMRI activity in the sensorimotor cortices modulated by the visual feedback, we defined 26 rectangular searchlights across sensorimotor cortices related to finger movement (see Methods). The activation was elevated in BA6 (PMd: dorsal premotor) in the continuous feedback condition. However, there was a reversal of the fMRI activity pattern between BA4a (anterior M1) and BA4p (posterior M1), with activation being higher in BA3a, 3b, 1, and 2 (S1) for the binary feedback condition. Indeed, ROI analysis found fMRI activity, which was higher in BA4a for the continuous feedback (*t*(23)=2.50, *p*<0.05) and higher in BA4p for the binary feedback (*t*(23)=2.34, *p*<0.05) (Fig. 2E). While the overall fMRI activity in the sensorimotor cortices was comparable (*F*(2, 46)=1.86, *p*=0.166), there was nearly significant difference between the feedback conditions (*F*(1, 23)=4.24, *p*=0.051). Importantly, we found a clear double dissociation of the fMRI activity between the feedback conditions across three representative ROIs, PMd, M1, and S1. The activity was higher in motor cortices (PMd and M1) for the continuous feedback (*t*(23)=3.68, *p*<0.01) and was higher in somatosensory cortices (S1) for the binary feedback (*t*(23)=4.38, *p*<0.001), resulting in significant interaction, effect between ROIs and feedback conditions (*F*(1, 23)=184.28, *p*<10^-4^).

Interestingly, a similar double dissociation of fMRI activity was also found in the visual cortices (Fig. 2B). While the overall activity across ROIs in the visual cortices (Early visual area, V3AB, and V7; *F*(1, 23)=3.45, *p*>0.05) was comparable, it was significantly different between the feedback conditions (*F*(1, 23) = 29.74, *p*<0.001). The early visual areas (V1, V2, and V3) were more activated when participants could not see the online cursor movement (*t*(23)=2.71, *p*<0.05); however, V3AB and V7 showed higher activation for the continuous feedback condition (*t*(23)=3.46, *p*<0.01; *t*(23)=11.98, *p*<0.001, respectively) (Fig. 2D), resulting highly significant interaction effect between ROIs and feedback conditions (*F*(1, 23)=97.60, *p*<0.001). Generally, early visual areas were more activated with the binary feedback, whereas higher-order visual areas were more activated with the continuous feedback.

### Connectivity between sensorimotor and visual cortices

Next, we examined whether the connectivity between sensorimotor and visual networks could explain *de novo* visuomotor skill learning. We employed task-based connectivity analysis due to its superiority over resting-state connectivity for the prediction of individual behaviors^16,17^. To complement the GLM analysis, we calculated a partial correlation controlling the effect of task-related coactivation^18^. In this way, we made the connectivity analysis orthogonal to the GLM analysis^19^.

We hypothesized that the connectivity would decrease as learning a hand-to-cursor mapping progressed with increasing autonomy of each network^1^. Furthermore, we sought to find which specific sensorimotor or visual regions are involved in *de novo* skill learning. To define nodes for connectivity analysis, we first chose the top five most significant clusters related to finger movement in the sensorimotor and visual cortices and then split the largest motor cortices into two regions, resulting in three regions for each sensorimotor (PMd, M1, and S1) and visual (dorsal, ventral, early) network (Fig. 3A). We focused on connectivity between sensorimotor network and visual networks.

**Figure 3.**
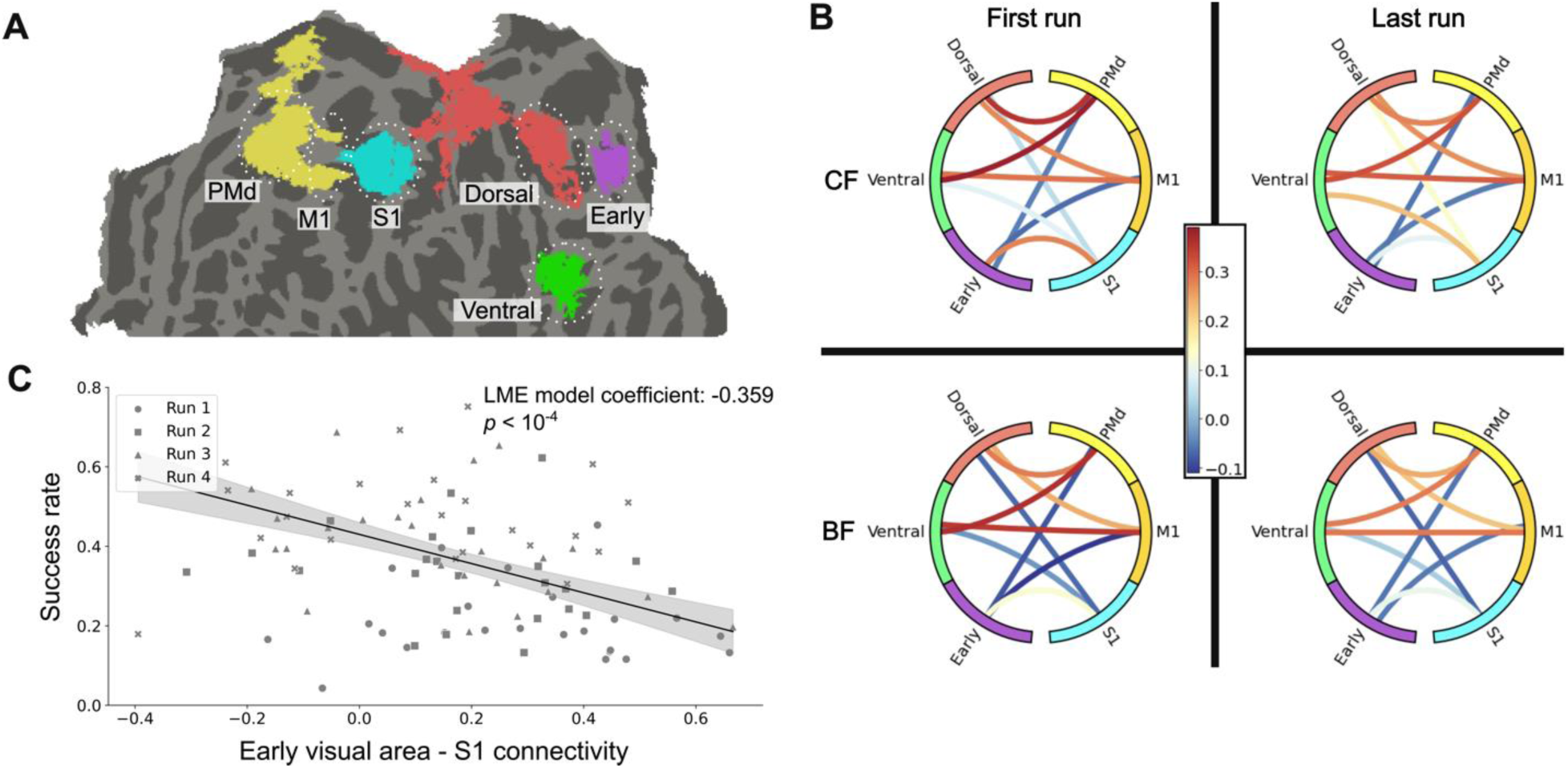
**(A)** Top five significant clusters from the GLM result. The yellow cluster was parcellated into PMd and M1. Only the posterior part of the violet cluster was used and defined as a dorsal visual region. **(B)** Functional connectivity across sensorimotor regions and visual regions. The color of the nodes is consistent with the color of the clusters in **A**. **(C)** The relationship between S1-early visual cortices connectivity and the success rate. A linear mixed-effect model was used to explain individual differences in motor skill learning. Each participant underwent four fMRI runs indicated by different shapes of dots.

Intriguingly, the connectivity patterns after controlling for task-specific effects were comparable with coactivation patterns under the two alternate conditions shown in Fig. 2 (Fig. 3B). In particular, we found higher connectivity between the motor cortices (PMd and M1) and the late visual regions (dorsal and ventral) than between the motor cortices and the early visual region regardless of conditions or fMRI runs (see Methods; CF and the first run: *t*(23)=10.80, *p*<10^-4^; CF and the last run: *t*(23)=10.33, *p*<10^-4^; BF and the first run *t*(23)=10.89, *p*<10^-4^; BF and the last run: *t*(23)=10.11, *p*<10^-4^; all Bonferroni-corrected). In contrast, the connectivity between S1 and the early visual region was higher than between S1 and the late visual regions in the first runs of both conditions (CF and the first run: *t*(23)=4.07, *p*<0.001; CF and the last run: *t*(23)=1.54, *p*=0.55; BF and the first run *t*(23)=3.93, *p*<0.001; BF and the last run: *t*(23)=2.54, *p*=0.07; all Bonferroni-corrected).

To investigate the relationship between the strength of visuomotor interaction and learning performance, we tested how connectivity between sensorimotor and visual cortices changes over time. Interestingly, among 18 tests for connectivity between three sensorimotor ROIs and three visual ROIs for each feedback condition, only the connectivity between S1 and early visual cortex significantly decreased from the first to the last fMRI run in the continuous feedback condition (*t*(23)=5.12, Bonferroni-corrected, *p*<0.001) (Fig. 3B). All the other connectivity did not reach significance level of corrected *p*<0.05 for multiple tests. Then, we correlated the connectivity reduction with the improvement in learning performance, measured as an averaged success rate during each feedback condition and fMRI run. A linear mixed effects model analysis (fixed effect: connectivity across four fMRI runs, random effect: participants) revealed that the reduction of connectivity between S1 and the early visual region was highly correlated with performance improvement when continuous feedback was provided (coef=-0.36, SE=0.077, *p*<0.001) (Fig. 3C).

## Discussion

In this work, we found distinctive involvement of the sensorimotor and visual cortices subregions in learning a motor skill *de novo*. We found clear double dissociation of fMRI activity in the subregions as well as their interaction, which were mediated by visual feedback. Motor and late visual cortices were more activated when continuous visual feedback was available than when only binary visual feedback was available. In contrast, somatosensory cortices and early visual cortex exhibited the opposite activation pattern. We also found a significant learning-induced reduction of connectivity between somatosensory and earl visual cortices, which is correlated with improved performance.

It is challenging to separate fMRI activity between closely located motor and somatosensory cortices since efferent motor commands and afferent proprioceptive feedback are processed simultaneously during continuous motor control^7^. In our experiment, manipulation of visual feedback allowed us to dissociate the fMRI activity by modulating the extent to which participants depended on visual or proprioceptive feedback to perform the task. When visual feedback is limited to binary feedback, participants are more likely to attend to proprioception, i.e., hand posture, leading to higher fMRI activity in somatosensory cortices than in motor cortices. That is, the extent of visual feedback switches attentional focus on the visual or proprioceptive modality as suggested in a previous study^11^. In our experiment of learning an arbitrary hand-to-cursor mapping, this dissociation would be more prominent than in typical motor learning tasks due to more considerable uncertainty about the motor and sensory goals of the task^7^. We found the boundary of the dissociation in the primary motor cortex. Specifically, the anterior part showed greater activation with the presence of online feedback compared to its absence, while the posterior part exhibited the reverse pattern. Our results are consistent with other studies claiming that the anterior M1 is more involved in motor execution, which is externally triggered, and the posterior M1 is more related to increased sensory attention^20,21^. Indeed, the anterior and posterior M1 were known to differ not only in their cytoarchitecture and neurochemistry but also in their functions^22–24^. Although our surface-based GLM analysis has the advantage over a volume-based GLM to delineate the boundary with less overlapped fMRI activity between M1 and S1^21^, the spatial resolution would not be enough to dissociate the functional difference. Future studies using 7 T fMRI with better spatial resolution and SNR would be necessary to dissociate the roles of sensorimotor subregions.

The dissociative responses observed in visual cortices could be explained by competition for cognitive resources between early and late visual cortices. In the online feedback condition, late visual cortices, including V5/MT, which are sensitive to visual motion, would be more activated than in early regions. On the other hand, in the binary feedback condition where no online visual feedback is provided, the cognitive resource would be released from the late visual cortices, resulting in higher activation in the early visual cortex^10,11^.

The task-based functional connectivity analysis further explains computational processes involved in *de novo* motor skill learning. Recent research has sparked debate around two contrasting motor control theories: the optimal control theory and the active inference theory^25^. A key point of contention lies in the active inference theory’s rejection of the concept of efference copy. Based on our results, the active inference theory would be a less plausible explanation for *de novo* motor learning since it assumes prior state estimation that participants do not have when learning a motor skill from scratch. In contrast, the optimal control theory would provide a better explanation for *de novo* motor learning as it suggests that M1 transmits an efference copy to the parietal cortex^26^, which is supported by our connectivity analysis results demonstrating a solid connection between M1 and the dorsal visual region in the parietal cortex. Furthermore, our primary finding of an apparent dissociation between M1 and S1 supports the optimal control theory, as it contradicts the active inference theory’s premise that action and perception cannot be separated^27^. Our claim aligns with an earlier study suggesting that S1 plays a role in receiving an efference copy as it encodes sensory information in anticipation of movements^28,29^. Future studies may be warranted to reconcile these two *de novo* motor learning explanations.

A landmark fMRI study in motor sequence learning revealed that visuomotor interaction is more dominant in an earlier stage of learning visuomotor mappings and decreases in the later stage of learning with more automatic performance^1^. Our results also found a significant reduction of visuomotor interaction, specifically between S1 and early visual cortex, during short-term *de novo* motor skill learning. The more significant contribution of S1 than M1 to learning is potentially due to the complexity of our task with highly uncertain sensory targets, which emphasizes the role of proprioception necessary to estimate sensory states (i.e., hand postures)^2,7^. On the other hand, the more significant contribution of the early instead of late visual cortices would be related to higher spatial selectivity of visual fields corresponding to target positions in the early visual cortex^30^. However, the current study needs to provide a clear explanation about the lack of contribution of the interaction between motor cortices and late visual cortices to motor learning. We speculate that the somatosensory cortex could play a more critical role in initial visuomotor learning when a sensory target is highly uncertain^2^, as in our task learning an arbitrary hand-to-cursor mapping. It would be intriguing to explore the contribution of S1 to *de novo* motor learning in a more advanced stage of learning with a longitudinal study^14^.

## Methods

### Subjects

Twenty-six healthy adults from the Sungkyunkwan University community participated in this study. According to the Edinburgh Handedness Inventory^31^, all participants were right-handed. In addition, they had no history of neurological or psychiatric disease and had normal or corrected-to-normal vision. Twenty-four participants (14 females; mean age = 24.9 ± 4.7 years, age range = 18-35 years) completed all the experiment sessions. Two participants dropped out in the middle of the experiment due to severe fatigue, and thus, their data were excluded from the analysis. All participants had corrected or corrected-to-normal vision and provided written consent. All the experimental procedures adhered to the Declaration of Helsinki and were approved by the Institutional Review Board of Sungkyunkwan University, Suwon, Republic of Korea (IRB No. 2018-05-003-032). Participants underwent two scanning sessions for 1.5 hours at a 3T fMRI scanner and received monetary rewards for their participation after the experiment.

### fMRI data collection

In the present study, fMRI data was acquired using a 3-T Siemens Magnetom Prisma scanner equipped with a 64-channel head coil. Functional images were obtained using an echo-planar imaging (EPI) sequence with specific parameters: a total of 300 volumes (with 310 volumes for localizer fMRI), a repetition time (TR) of 2,000 ms, an echo time (TE) of 35.0 ms, a flip angle (FA) of 90°, a field of view (FOV) measuring 200 mm, a matrix size of 101 X 113 X 91 voxels, 72 axial slices, and a slice thickness of 2.0 mm. For anatomical referencing, a T1-weighted anatomical scan of the entire brain was conducted using a magnetization-prepared rapid acquisition with gradient echo MPRAGE sequence. This anatomical scan employed specific parameters, including a TR of 2,300 ms, a TE of 2.28 ms, an FA of 8°, an FOV of 256 mm, a matrix size of 204 X 262 X 260 voxels; 192 axial slices, and a slice thickness of 1.00 mm. Before the functional scans, two EPI images were acquired with opposite-phase encoding directions (posterior-to-anterior and anterior-to-posterior) to enable subsequent distortion correction.

### Data glove

Participants wore the MR-compatible data glove (5DT Glove 14 Ultra) on their right hand. With the data glove, hand postures recorded by 14 sensors were converted to cursor positions on the 2-dimensional screen. The hand-to-cursor mapping is defined below.

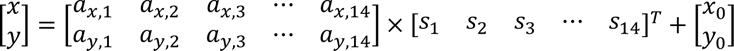

where 𝑠_𝑘_(𝑘 = 1,2, …, 14) indicates each of the 14 sensor inputs from the data glove, and 𝑥 and 𝑦 indicate the horizontal and vertical position of the cursor. The time-series data of the 14 sensors was sampled at 60 Hz. The above equation can be rewritten as 𝐫 = 𝐀𝐬 + 𝐫_𝟎_, where the mapping matrix 𝐀 and the offset 𝐫_𝟎_ were determined from the calibration and localizer session.

### Calibration

Before fMRI scanning, participants had a calibration phase in an MRI scanning control room. After wearing the data glove, participants moved their right hands while they could see a graph of 14 bars moving in real time based on the 14 sensors. Participants were instructed to try various combinations of hand postures. Next, we conducted a principal component analysis with the covariance matrix derived from the time series collected from the 14 sensors. The first two principal components were employed to create the mapping matrix 𝐀, and the offset 𝐫_𝟎_ was calculated to ensure that the average hand posture was aligned with the center of the screen. After that, we ensured that participants could visit all 25 cells of a 5×5 grid (Fig. 1A).

### Localizer session

After the calibration phase, participants moved to an MRI scanning room. They lay on their back in the scanner, observing the screen via a mirror. Foam pads were applied to all participants to reduce head movement. In addition, they wore the data glove on their right hand and put the hand in a comfortable position. Notably, participants could not see their hands moving during the entire fMRI experiment.

To localize finger movement regions in the brain, participants had a localizer session before the main task session. When “Move” was displayed on the screen, participants performed natural-speed movements with their right fingers, ceasing their movements upon the appearance of the “Stop” text. Each “Move” or “Stop” condition had a duration of 48 seconds, separated by 2-second intervals, and a total of six sets of “Move” and “Stop” conditions were executed. To make sure it is de novo learning to participants, we recalibrated the mapping matrix 𝐀 and the offset 𝐫_𝟎_ using the data acquired from the finger movements in the last two “Move” blocks. We also ensured that all 25 grid cells were reachable by finger movements.

### Main task session

Participants were required to move their right fingers to control a cursor and reach a target cell. A target cell is a gray cell with a yellow crosshair in its center. If participants reached a target cell, its color changed to red, and they were required to stay in the target cell until the next one appeared. In other words, the goal of the main task for participants was to reach a target as quickly as possible and to stay as long as possible in the target. A target cell appeared for 4 s. Regardless of whether participants reached a target or not, the target was changed to the next one after 4 s. Each block had 12 trials with a sequence of either 13-3-25-21-13-25-3-21-25-13-21-3 (Sequence 1: triangle) or 13-23-5-1-13-5-23-1-5-13-1-23 (Sequence 2: inverted triangle), where the number indicates the grid cell number determined by the formula 𝑘 = 5𝑖 + 𝑗 − 5, with 𝑖 representing the row index and 𝑗 representing the column index. Half of the participants were presented with Sequence 1, and the other half were presented with Sequence 2 for counterbalancing. In total, there were four runs, and each run consisted of 12 blocks. Thus, the duration of each run was about 576 s (144 trials × 4 s).

Notably, there were two experimental conditions: full visual feedback and binary visual feedback. In the full visual feedback condition, participants could see both cursor movement and targets. On the other hand, in the binary visual feedback condition, participants could only see targets. However, the cursor was not visible (Fig. 1B). Thus, participants could only guess their cursor position by the color of the targets. The two conditions were applied in separate blocks alternately. During odd-numbered blocks, the cursor position was continuously represented by a white crosshair, whereas it was hidden during even-numbered blocks.

### Behavioral data analysis

The proportion of time during which a target turned red was measured as a trial-by-trial success rate. For example, the success rate of one block was defined by the amount of time targets turned red during one block divided by 48 s, a total time of one block. We used a two-sided paired t-test between different feedback conditions, or fMRI runs to see if there was a significant difference. MATLAB (version R2022a, MathWorks), Python (version 3.10.8), and Jupyter notebook (version 6.4.12) were utilized for all statistical analysis.

### fMRI data analysis: preprocessing

MRI data were analyzed with FreeSurfer (version 7.2.0)^32^. We first followed the FreeSurfer ‘recon-all’ pipeline for the preprocessing of anatomical data. Then, for functional data, we followed the FreeSurfer Functional Analysis Stream (FS-FAST) preprocessing pipeline (https://surfer.nmr.mgh.harvard.edu/fswiki/FsFastTutorialV6.0/FsFastPreProc). Functional data from each subject were registered to the same-subject FreeSurfer anatomical data with motion correction and slice-timing correction. Then, the functional data were resampled to common FreeSurfer space. Note that we did not perform spatial smoothing because we wanted to investigate precise brain mapping to distinguish sensorimotor areas. The non-smoothed data were also used for connectivity analysis.

### fMRI data analysis: GLM

We utilized FS-FAST for the GLM analysis^33^. We utilized FS-FAST for the GLM analysis. To identify regions responding to full visual feedback and only binary visual feedback, we used a blocked task design for the regressors of interest. Then, the regressors of interest were convolved with the SPM canonical hemodynamic response function (HRF) of zero derivatives. It was not smoothed here as well. For nuisance regressors, we used motion parameters created in preprocessing. The top three components were used. In addition, third-order polynomial regressors were included. After the first-level analysis, individual results were concatenated into one file with the “isxconcat-sess” function. Then, group GLM was performed with “mri_glmfit .” Multiple comparisons correction was done by “mri_glmfit-sim” with cluster-forming threshold (CFT) of *p*<10^-4^ and cluster-wise p-value of *p*<0.05. The final full width at half maximum (FWHM) of the inherent smoothness was taken into account for multiple comparisons correction. A similar analysis was also done on the localizer result with different conditions of “Move” and “Stop” to identify regions responding to hand movement.

### ROI selection

We defined ROIs in PMd, M1, S1, V1, V2, V3, V3A, V3B and V7 based on the GLM result. ROIs in sensorimotor areas were defined as an overlap between clusters obtained from the localizer result and Brodmann areas provided by FreeSurfer30 (BA6 for PMd; BA4a and 4p for M1; BA1, 2, 3a and 3b for S1). For visual areas, we used the surface-based atlas made by Wang, et al31. They grouped V1d, V1v, V2d, V2v, V3d, and V3v regions, defining them as early visual region, and we adhered to their classification. For precise brain mappings, we subdivided sensorimotor areas spanning PMd, M1, and S1. Using FreeSurfer’s fsaverage brain, we cut the ROIs at regular distances as parallel as possible to the boundaries between the ROIs (Fig. 2C). These ROIs were employed in the univariate activation analysis, where beta estimates obtained from the GLM analysis within the ROIs are averaged (Fig. 2C, 2E).

For connectivity analysis, we chose the five most significant clusters made from the GLM result (Fig. 3A). On the other hand, based on our hypotheses, we divided the yellow cluster into PMd and M1 parts. We also used only the posterior part of the scarlet cluster and defined it as a dorsal visual cluster. As a result, we have six clusters of PMd, M1, S1, dorsal, ventral, and early visual regions. Then, we defined individual circular masks of 3mm radius from each cluster. First, using “mri_surfcluster”, we identified each vertex number, the center of a circular mask, with the highest beta estimates. After that, the circular masks were drawn around the center with the “mri_volsynth”, “mris_fwhm”, and “mri_binarize” functions.

### fMRI data analysis: connectivity

The task-based time-series data within the seeds defined in ROIs were averaged and extracted by “mri_segstats”. For the time-series data of each run, we regressed out the same motion parameters and third-order polynomials as used in the GLM analysis, as well as other nuisance variables such as global signal, white matter (WM), ventricles, and cerebrospinal fluid (VCSF). Both VM and VCSF regressors were defined by the top component from PCA by “fcseed-sess” respectively. To remove task-specific effects, functional connectivity between two seeds was calculated by partial correlation with covariances of both conditions from the design matrix used in the GLM analysis. The functional connectivity was calculated between the sensorimotor regions (PMd, M1, and S1) and the visual regions (early, dorsal, and ventral).

At first, for direct comparison with the GLM result that the motor cortices and the late visual regions coactivated under the continuous feedback condition, and S1 and the early visual region coactivated under the binary feedback condition, we averaged connectivity values of the motor nodes and the late visual nodes. For instance, when we performed a two-tailed paired t-test with S1-early and S1-late connectivity, we averaged connectivity between S1-dorsal and S1-ventral.

Next, to check whether there was a significant difference in functional connectivity between the sensorimotor regions and the visual regions across the first and the last runs, we performed a two-tailed paired t-test. 18 separate t-tests (three sensorimotor regions × three visual regions × two conditions) were conducted, and Bonferroni-corrected p-value was applied to see the difference. Following the observation of a significant decrease in functional connectivity between S1 and the early visual region, we further conducted the linear mixed effects (LME) model analysis with random intercepts (fixed effect: connectivity across four fMRI runs, random effect: participants). The purpose of the LME analysis was to examine the correlation between the observed changes in functional connectivity and behavioral performance improvement.

## Code and Data Availability

All MRI data used in this study are archived in the Hanyang University Network Attached Storage (NAS). The corresponding author can provide any information on the dataset and codes necessary to generate results and figures shown in this study.

## Acknowledgments

Neuroimaging was performed at the Center for Neuroscience Imaging Research located at Sungkyunkwan University, supported by the Institute for Basic Science.

## Declaration of generative AI and AI-assisted technologies in the writing process

During the preparation of this work, the authors used ChatGPT4.0 and Grammarly to edit the manuscript with grammar checks. After using these tools, the authors reviewed and edited the content as needed and took full responsibility for the content of the publication.

